# Insecticidal Effects of Datura Species against Major Agricultural Pests

**DOI:** 10.1101/2023.10.28.564503

**Authors:** Meenakshi Sharma, Vikas Kumar Singh, Deepti Chaturvedi, Anil Kumar Delta, Madan Mohan Sharma, Prashant Kaushik

## Abstract

Insecticides derived from plants provide environmentally friendly and sustainable options as alternatives to synthetic chemicals. This investigation delves into the insecticidal capabilities of two distinct Datura species, namely, Datura alba and Datura stramonium, in combatting significant pests such as *Spodoptera litura, Bemesia tabaci*, and *Callosobruchus maculatus*. Datura plants have a well-documented history of medicinal use and are recognized for their abundant Tropane alkaloid content, which acts as an innate defense mechanism against pests. Hence, the current investigation involved the extraction of chemical compounds from the leaves, flowers, seeds, and roots of both Datura stramonium and Datura alba. Subsequently, these extracts were assessed for efficacy using Spodoptera litura larvae*, Bemesia tabaci*, and *Callosobruchus maculatus* in bioassays. The larvicidal experiments encompassed a range of concentrations (3%, 5%, 10%), with larval mortality being documented at multiple time intervals: 6, 12, 24, 48, and 72 hours. The findings from our investigation uncover the notable insecticidal attributes of root, leaf, flower, and seed extracts from both *Datura alba* and *Datura stramonium* against these larvae. These extracts demonstrated distinct mortality rates at various concentrations and exposure durations. This comprehensive analysis imparts crucial knowledge regarding the insecticidal qualities of Datura species and their prospective contributions to sustainable pest control practices. The observed effectiveness of these plant extracts against a spectrum of significant pests hints at their potential as ecological alternatives within integrated pest management strategies.

## 1. Introduction

Advanced plant species have increasingly become a valuable reservoir for pioneering insecticides (Dev and Koul 1997). Plants possess the capability to produce natural compounds as a defense mechanism against diseases, pathogens, and specific substances with pesticidal properties (Benhamou et al., 2012; European Commission, 2020). For generations, traditional practices worldwide have harnessed the insect-repelling qualities of plant materials (Belmain et al., 2001). In contrast to synthetic options, botanical insecticides present promising benefits such as environmental friendliness, reduced expenses, straightforward processing, and applicability for farmers and small industries. The Datura genus, a member of the Solanaceae family, has a rich history of medicinal utilization, including in contemporary India (Priya et al., 2002). Currently, China holds the dominant global position in producing and exporting medicinal plants, with India closely trailing as a significant player (Singh and Kumar, 2021).

Referred to as Thorn apple plants, these herbaceous or short-lived perennial plants find extensive use across various regions, including the Mediterranean and tropical and subtropical areas worldwide. A distinctive feature common to all members of the Datura genus is their significant abundance of Tropane alkaloids, which represent a class of secondary metabolites. These alkaloids are produced in the roots, stored within vacuoles, and released as a chemical defense against pests (Evans, 1979; Conklin, 1976). Botanical insecticides generally demonstrate selective effectiveness against a narrow spectrum of target species and tend to break down into non-toxic substances as they biodegrade. Kim et al. (2003) noted that this attribute, when paired with the potential to include plants in integrated pest management strategies, enables the creation of new, safer insect control agents. Moreover, many plant species, especially those in tropical regions, may serve as botanical insecticides or as sources of bioactive compounds, as acknowledged by Saxena et al. (1992) and Shaalan et al. (2005).

*Datura stramonium* is a tall, annual, and branching flowering plant that originally hails from the Americas but is now widespread globally, including Europe, Central and North America, South America, Asia, and Africa (Bayih, 2014). *Datura stramonium* is an upright, malodorous herb with a bushy growth pattern, reaching heights ranging from 2 to 5 feet. It boasts a lengthy, sturdy, fibrous, and pale-white root. The stem is cylindrical, upright, devoid of hair, and may exist in a single or forked form. The leaves are sleek, serrated, and velvety, featuring an irregularly wavy outline and retaining a bitter and nauseating flavor, which remains even after drying. The fruit measures approximately 5cm in length and comprises four-valve capsules densely adorned with thorns, reminiscent of a walnut size. Upon reaching maturity, the fruit splits into four chambers, each housing numerous seeds. These seeds are elongated, flat, kidney-shaped, and have a black coloration (Soni, 2012; Li, 2012; Usha et al., 2009; Olofintoye, 2011; Tostes, 2002).

*Datura alba* Nees, a medicinal plant found in warmer regions across the globe, is commonly referred to as Angel’s trumpet or Devil’s trumpet and typically reaches an average height of 1.5 meters (Iyekowa et al., 2023). This plant exhibits tolerance to heavy metals, thriving in locations contaminated with these substances (Fang et al. 2013). Earlier studies have explored essential oils’ efficacy for fumigating adult insects and their larvae. Recent investigations have focused on examining the toxicity of essential oils, including their major constituents, through contact and fumigation methods against insect eggs that infest stored products (Shaaya et al. 1993; Ho et al. 1997; Huang et al. 1997, 2000). These compounds represent valuable reservoirs of bioactive agents for both medical applications and pest management (Bourgaud et al. 2001). Datura plants are notably rich in scopolamine (Shonle and Bergelson, 2000; Gerlach, 2006), which has demonstrated adverse effects on various insects (Hsiao and Fraenkel, 1968; Krug and Proksch, 1993).

Beyond their relevance in the pharmaceutical field, the quantification and study of these compounds hold potential importance in agricultural contexts. For instance, fluctuations in their levels may offer insights into the plant’s general well-being, its strategies for pest resilience, and its capacity to acclimate to particular environmental circumstances (Jabeen et al., 2022). Moreover, studies have shown these plants can combat agricultural insect pests, including onion thrips (Malik, 2005), cabbage butterflies, rice ear-cutting caterpillars (Zhou et al., 2008), and aphids, grasshoppers, and ants (Kuganathan et al., 2008; Kuganathan & Ganeshalingam, 2011). Researchers have demonstrated their efficacy against storage beetles, grain borers, and flour beetles (Tierto Niber et al., 1992; Pascual-Villalobos & Robledo, 1998; Dwivedi & Shekhawat, 2004; Kamruzaman et al., 2005).

The active compounds extracted from Datura plants show promise for inclusion in pest control initiatives, especially for managing grain storage pests. Within this context, the objective of this research is to present the findings regarding the toxicity of two Datura species (*Datura alba* and *Datura stramonium*) against three significant pests: *Spodoptera litura*, *Bemesia tabaci*, and *Callosobruchus maculatus*.

## 2. Materials and Methods

### 2.1 Insect Culture

The larvae of *Spodoptera litura*, *Bemesia tabaci* and *Callosobruchus maculatus* were obtained from insect cultures maintained at the Entomology Department Research Lab, located in Jharkhand, India. The insects were reared under controlled conditions (25±2°C, 65±5% RH, 14:10 h L:D photoperiod) as described by (Hu et al. 2019). Newly hatched larvae were used for the bioassays.

### 2.2. Plant Collection, Identification and Extraction

The leaves, flowers, seeds, and roots of *Datura stramonium* and *Datura alba* were obtained from the medicinal garden of the Department of Botany, Kurukshetra University, Kurukshetra, India. The plant materials were shade-dried, powdered mechanically, and stored in airtight containers. For extraction, 200 g of each powdered plant part was extracted with 95% ethanol using the Soxhlet apparatus for 18 hours. The extracts were filtered and concentrated under reduced pressure in a rotary evaporator at 50°C. The concentrated extracts were stored at 4°C until testing.

### 2.3 Larvicidal Bioassay

The larvicidal activity was evaluated following standard protocol (Harve et al. 2004; Mallick et al. 2015). Different concentrations (3%, 5%, 10%) of each plant extract were prepared in acetone. Twenty larvae were placed in Petri dishes lined with filter paper. The extracts were applied topically on the larvae using a micropipette. Controls received the solvent only. Larval mortality was recorded after 6, 12, 24, 48 and 72 hours. Mortality was corrected using Abbott’s formula (Ali et al. 2020). The bioassays had 3 replicates and were repeated 2 times.

### 2.4. Statistical analysis of phenotypic data

Larval mortality percentages were calculated using the formula: Mortality % = (number of dead larvae / number of introduced larvae) × 100. To eliminate environmental effects, the best linear unbiased predictor (BLUP) of the calculated larval percentages was estimated using the Meta-R program V.6.0 (Alvarado et al., 2017). The R program (Team RC, 2016) with various packages (Openxlsx, lme4, emmeans, agricolae, reshape, reshape2, car, metan, multcompView) was used for analysis of variance (ANOVA), treating plants, plant parts, concentrations, replications, pathogens, and HPI as fixed and random effects. Statistical significance was determined at P < 0.001, and results were evaluated using the least significant difference (LSD) method at p < 0.05 with R packages tidyverse, ggthemes, multcompView, dplyr, ggplot2.

### 2.5. Determining the LC50 values

The probit analysis procedure follows the method described by Finney (1971) and was performed using GW-BASIC software (1985). This involves maximum likelihood estimation to fit a straight line to the probit-transformed mortality data as a function of logarithmic dose. The slope and intercept parameters of the line are iteratively adjusted to obtain a best fit model. The LC50 value is determined from the fitted regression line as the concentration corresponding to 50% mortality after back-transforming from the probit scale. The 95% confidence limits for the LC50 were also calculated from the fitted model. The LC50 values for different plant extracts were compared using these 95% confidence intervals. Non-overlapping confidence intervals indicate statistically significant differences between the LC50 values at the 5% level. This allows assessment of the relative potency of the various plant extracts based on the concentration required to induce 50% larval mortality. The probit analysis provides a robust statistical method to quantify the relationship between larval mortality and extract dose. Determining the LC50 values along with 95% confidence intervals enables clear comparison of the effectiveness of different plant extracts against insect pests.

## 3. Results

Larval mortalities obtained from bioassays of extracts of *Dhatura stramonium*, and *Dhatura alba*, roots, leaves, flowers, and seeds against *Spodoptera litura, Bemesia tabaci,* and *Dhatura stramonium* larvae after 6 hours, 12 hours, 24 hours, 48 hours, and 72 hours exposure periods. All plant extracts showed low, moderate, and high larvicidal activities tested between 3 to 10% concentration (Table 1).

This study evaluated the insecticidal properties of *Dhatura stramonium* plant extracts derived from various plant parts (root, leaf, flower, and seed) against *Spodoptera litura* larvae. The experiment assessed the mortality rates of the larvae at different concentrations (3%, 5%, and 10%) and over five time intervals (6, 12, 24, 48, and 72 hours). The results demonstrated that the mortality percentage of *Spodoptera litura* larvae was concentration and time-dependent. Higher concentrations are generally correlated with increased larval mortality. For example, at a 10% concentration, the mortality rate ranged from 25.00% at 24 hours to 76.25% at 72 hours. The efficacy of the extracts varied across plant parts and concentrations. For instance, the leaf extract at 10% concentration led to a mortality rate of 52.50% at 72 hours (Table 1).

**Table 1:** Mortality percentage of *Dhatura stramonium* plant’s root, leaf, flower, and seed extracts against *Spodoptera litura*.

| Plant part | Concentration (%) | 6 h | 12 h | 24 h | 48 h | 72 h |
| --- | --- | --- | --- | --- | --- | --- |
| Root | 3 | 3.75 | 5.00 | 7.50 | 15.00 | 21.25 |
|  | 5 | 5.00 | 6.25 | 13.75 | 21.25 | 25.00 |
|  | 10 | 12.50 | 20.00 | 25.00 | 31.25 | 38.75 |
| Leaf | 3 | 5.00 | 8.75 | 13.75 | 21.25 | 25.00 |
|  | 5 | 5.00 | 10.00 | 18.75 | 25.00 | 30.00 |
|  | 10 | 17.50 | 25.00 | 33.75 | 41.25 | 52.50 |
| Flower | 3 | 5.00 | 5.00 | 10.00 | 16.25 | 21.25 |
|  | 5 | 5.00 | 8.75 | 16.25 | 22.50 | 27.50 |
|  | 10 | 15.00 | 20.00 | 27.50 | 33.75 | 40.00 |
| Seed | 3 | 7.50 | 11.25 | 18.75 | 25.00 | 30.00 |
|  | 5 | 8.75 | 16.25 | 25.00 | 33.75 | 42.50 |
|  | 10 | 21.25 | 37.50 | 57.50 | 67.50 | 76.25 |

The mortality rates of *Bemesia tabaci* larvae exposed to extracts obtained from various parts of *Dhatura stramonium* plants, including the root, leaf, flower, and seed at the same time point and concentration. Notably, higher concentrations were associated with increased mortality. For example, at a 10% concentration, the mortality rate ranged from 33.75% at 24 hours to 67.50% at 72 hours for the root extract. Furthermore, the efficacy of the extracts displayed variations based on the plant part and concentration. As an illustration, the leaf extract at 10% concentration resulted in a mortality rate of 60.00% at 72 hours (Table 2).

**Table 2:** Insecticidal effects of *Dhatura stramonium* plants root, leaf, flower, and seed extracts against *Bemesia tabaci*.

| Plant part | Concentration (%) | 6 h | 12 h | 24 h | 48 h | 72 h |
| --- | --- | --- | --- | --- | --- | --- |
| Root | 3 | 5.00 | 10.00 | 16.25 | 25.00 | 31.25 |
|  | 5 | 10.00 | 18.75 | 30.00 | 40.00 | 46.25 |
|  | 10 | 13.75 | 20.00 | 33.75 | 43.75 | 48.75 |
| Leaf | 3 | 6.25 | 15.00 | 23.75 | 31.25 | 43.75 |
|  | 5 | 11.25 | 25.00 | 38.75 | 47.50 | 56.25 |
|  | 10 | 18.75 | 25.00 | 38.75 | 52.50 | 60.00 |
| Flower | 3 | 5.00 | 11.25 | 18.75 | 28.75 | 35.00 |
|  | 5 | 11.25 | 21.25 | 32.50 | 41.25 | 50.00 |
|  | 10 | 15.00 | 23.75 | 38.75 | 46.25 | 53.75 |
| Seed | 3 | 10.00 | 20.00 | 35.00 | 45.00 | 52.50 |
|  | 5 | 16.25 | 31.25 | 45.00 | 56.25 | 65.00 |
|  | 10 | 20.00 | 38.75 | 53.75 | 62.50 | 67.50 |

Similarly, this study assessed the insecticidal potential of *Dhatura stramonium* plant extracts from the root, leaf, flower, and seed against *Callosobruchus maculatus* at the similar conditions. Similar to the above, results demonstrated concentration-dependent mortality, with higher concentrations yielding increased mortality rates. For example, at 10% concentration, root extract resulted in mortality rates from 33.75% at 24 hours to 53.75% at 72 hours. Extract efficacy varied by plant part and concentration, e.g., 10% leaf extract led to 61.25% mortality at 72 hours. Extended exposure, particularly at 72 hours, consistently increased mortality (Table 3).

**Table 3:**
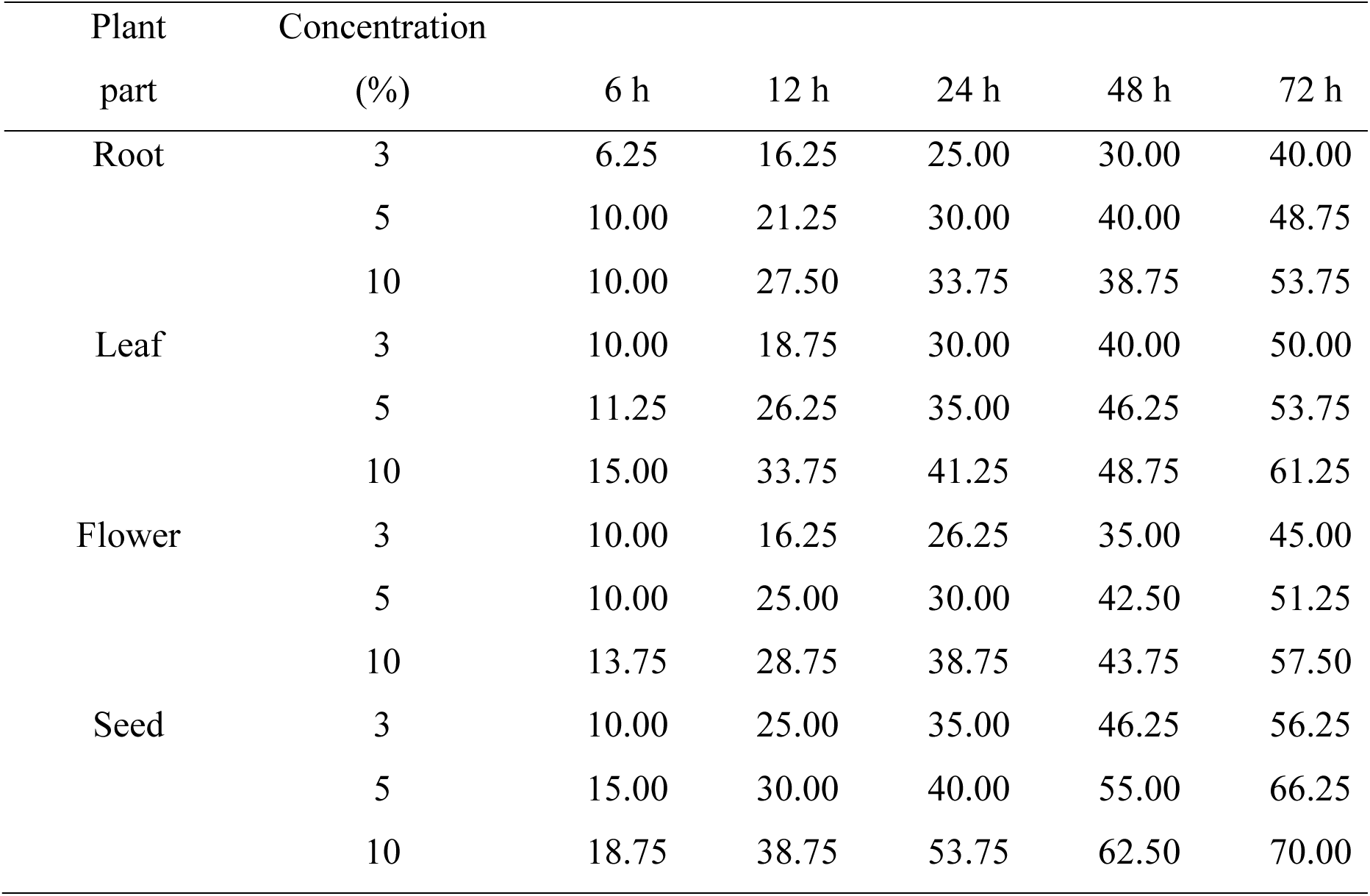
Insecticidal effects of *Dhatura stramonium* plants root, leaf, flower, and seed extracts against *Callosobruchus maculatus*.

| Plant part | Concentration (%) | 6 h | 12 h | 24 h | 48 h | 72 h |
| --- | --- | --- | --- | --- | --- | --- |
| Root | 3 | 6.25 | 16.25 | 25.00 | 30.00 | 40.00 |
|  | 5 | 10.00 | 21.25 | 30.00 | 40.00 | 48.75 |
|  | 10 | 10.00 | 27.50 | 33.75 | 38.75 | 53.75 |
| Leaf | 3 | 10.00 | 18.75 | 30.00 | 40.00 | 50.00 |
|  | 5 | 11.25 | 26.25 | 35.00 | 46.25 | 53.75 |
|  | 10 | 15.00 | 33.75 | 41.25 | 48.75 | 61.25 |
| Flower | 3 | 10.00 | 16.25 | 26.25 | 35.00 | 45.00 |
|  | 5 | 10.00 | 25.00 | 30.00 | 42.50 | 51.25 |
|  | 10 | 13.75 | 28.75 | 38.75 | 43.75 | 57.50 |
| Seed | 3 | 10.00 | 25.00 | 35.00 | 46.25 | 56.25 |
|  | 5 | 15.00 | 30.00 | 40.00 | 55.00 | 66.25 |
|  | 10 | 18.75 | 38.75 | 53.75 | 62.50 | 70.00 |

This study assessed the insecticidal properties of *Dhatura alba* plant extracts from the root, leaf, flower, and seed against *Spodoptera litura* larvae. Concentrations of 3%, 5%, and 10% were applied over 6 to 72 hours. The results demonstrated that higher concentrations consistently led to increased larval mortality. For example, at a 10% concentration, the root extract resulted in mortality rates ranging from 16.25% at 24 hours to 21.25% at 72 hours. The efficacy of the extracts varied across plant parts and concentrations, with the leaf extract at 10% concentration, resulting in a 40.00% mortality rate at 72 hours (Table 4).

**Table 4:** Insecticidal effects of *Dhatura alba* plants root, leaf, flower, and seed extracts against *Callosobruchus maculatus*.

| Plant part | Concentration (%) | 6 h | 12 h | 24 h | 48 h | 72 h |
| --- | --- | --- | --- | --- | --- | --- |
| Root | 3 | 0.00 | 2.50 | 6.25 | 8.75 | 12.50 |
|  | 5 | 1.25 | 5.00 | 8.75 | 11.25 | 16.25 |
|  | 10 | 6.25 | 11.25 | 16.25 | 21.25 | 21.25 |
| Leaf | 3 | 1.25 | 5.00 | 11.25 | 13.75 | 18.75 |
|  | 5 | 2.50 | 8.75 | 13.75 | 17.50 | 22.50 |
|  | 10 | 7.50 | 15.00 | 21.25 | 31.25 | 40.00 |
| Flower | 3 | 1.25 | 5.00 | 8.75 | 12.50 | 17.50 |
|  | 5 | 3.75 | 5.00 | 10.00 | 13.75 | 18.75 |
|  | 10 | 7.50 | 12.50 | 18.75 | 26.25 | 33.75 |
| Seed | 3 | 3.75 | 7.50 | 15.00 | 20.00 | 26.25 |
|  | 5 | 6.25 | 10.00 | 16.25 | 25.00 | 31.25 |
|  | 10 | 12.50 | 22.50 | 38.75 | 52.50 | 62.50 |

This study evaluated the insecticidal potential of *Dhatura alba* plant extracts from the root, leaf, flower, and seed against *Bemesia tabaci* over different concentrations (3%, 5%, and 10%) and time intervals (6 to 72 hours). Concentration and time dependence were observed, with higher concentrations resulting in increased mortality. For instance, at a 10% concentration, the root extract led to mortality rates from 25.00% at 24 hours to 42.50% at 72 hours. Efficacy varied by plant part and concentration, such as the leaf extract at 10% concentration resulting in a 51.25% mortality rate at 72 hours. Longer exposure times they have consistently produced higher mortality rates (Table 5).

**Table 5:** Insecticidal effects of *Dhatura alba* plants root, leaf, flower, and seed extracts against *Bemesia tabaci*.

| Plant part | Concentration |  |  |  |  |  |
| --- | --- | --- | --- | --- | --- | --- |
|  | (%) | 6 h | 12 h | 24 h | 48 h | 72 h |
| Root | 3 | 1.25 | 5.00 | 11.25 | 17.50 | 21.25 |
|  | 5 | 6.25 | 12.50 | 18.75 | 25.00 | 30.00 |
|  | 10 | 10.00 | 15.00 | 25.00 | 36.25 | 42.50 |
| Leaf | 3 | 3.75 | 11.25 | 17.50 | 25.00 | 36.25 |
|  | 5 | 6.25 | 13.75 | 22.50 | 32.50 | 42.50 |
|  | 10 | 13.75 | 20.00 | 30.00 | 41.25 | 51.25 |
| Flower | 3 | 3.75 | 10.00 | 15.00 | 23.75 | 27.50 |
|  | 5 | 8.75 | 15.00 | 27.50 | 32.50 | 40.00 |
|  | 10 | 8.75 | 16.25 | 25.00 | 35.00 | 47.50 |
| Seed | 3 | 7.50 | 15.00 | 26.25 | 32.50 | 45.00 |
|  | 5 | 13.75 | 23.75 | 36.25 | 43.75 | 58.75 |
|  | 10 | 12.50 | 26.25 | 38.75 | 51.25 | 60.00 |

This study assessed the insecticidal potential of *Dhatura alba* plant extracts (root, leaf, flower, seed) against *Callosobruchus maculatus* at different concentrations (3%, 5%, 10%) and time intervals (6 to 72 hours). Concentration-dependent mortality was observed, with higher concentrations leading to increased death rates. For instance, at 10% concentration, the root extract resulted in a mortality rate ranging from 26.25% at 24 hours to 45.00% at 72 hours. Efficacy varied by plant part and concentration, such as the leaf extract at 10% concentration leading to a 55.00% mortality rate at 72 hours. Longer exposure times they have consistently produced higher mortality rates (Table 6).

**Table 6:** Insecticidal effects of *Dhatura alba* plants root, leaf, flower, and seed extracts against *Callosobruchus maculatus*.

| Plant part | Concentration (%) | 6 h | 12 h | 24 h | 48 h | 72 h |
| --- | --- | --- | --- | --- | --- | --- |
| Root | 3 | 2.50 | 8.75 | 15.00 | 23.75 | 28.75 |
|  | 5 | 6.25 | 12.50 | 22.50 | 31.25 | 38.75 |
|  | 10 | 7.50 | 18.75 | 26.25 | 33.75 | 45.00 |
| Leaf | 3 | 5.00 | 15.00 | 25.00 | 30.00 | 41.25 |
|  | 5 | 10.00 | 18.75 | 27.50 | 36.25 | 50.00 |
|  | 10 | 12.50 | 22.50 | 32.50 | 41.25 | 55.00 |
| Flower | 3 | 5.00 | 13.75 | 22.50 | 30.00 | 38.75 |
|  | 5 | 7.50 | 18.75 | 28.75 | 35.00 | 47.50 |
|  | 10 | 10.00 | 20.00 | 28.75 | 37.50 | 51.25 |
| Seed | 3 | 6.25 | 13.75 | 23.75 | 28.75 | 41.25 |
|  | 5 | 8.75 | 17.50 | 28.75 | 42.50 | 58.75 |
|  | 10 | 12.50 | 25.00 | 40.00 | 50.00 | 61.25 |

The above results indicate a potential influence of the source material on insecticidal activity. Longer exposure times, particularly at 72 hours, led to elevated mortality rates. This data provides valuable insights into the insecticidal potential of *Dhatura* plant extracts against all three insects as mentioned earlier and offers guidance for pest management and agricultural application.

### 3.1. Statistical analysis of phenotypic data

The results of the ANOVA for all studied traits are presented in Table 7, 8. It is apparent that variations due to plants, plant part, concentration, replication, pathogens, HPI and its interactions like, plants: concentration, plants:pathogen, plants:part, and plants:HPI were all highly significant. This is expected due to complex phenotyping.

**Table 7:**
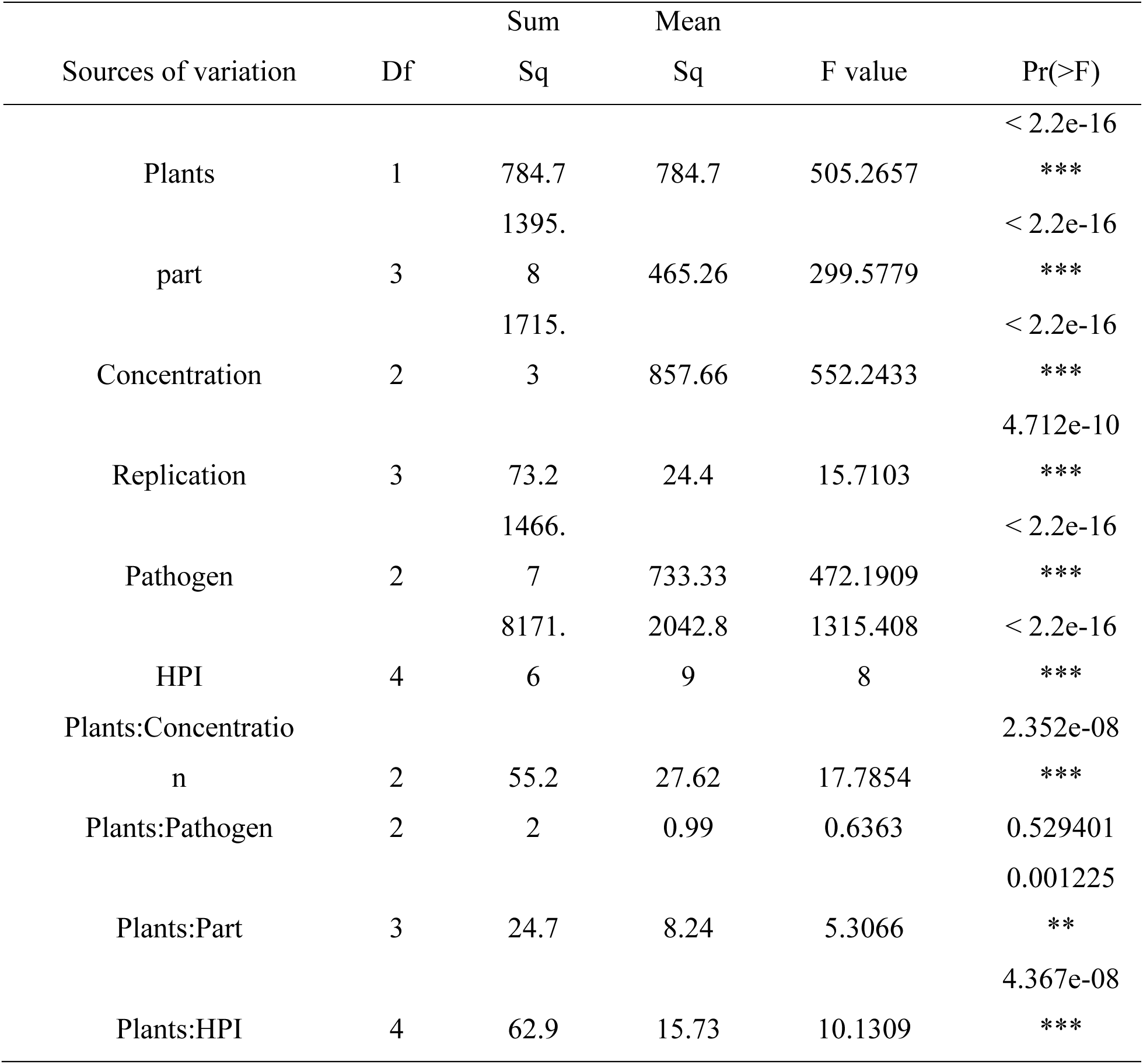
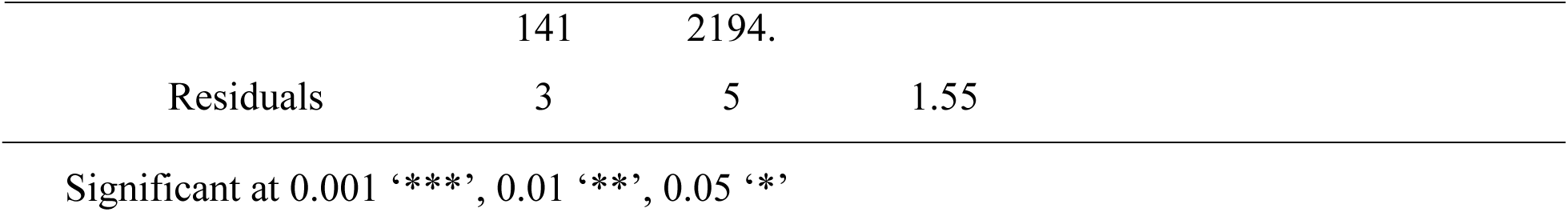
Analysis of variance of No. of mortality.

**Table 8:**
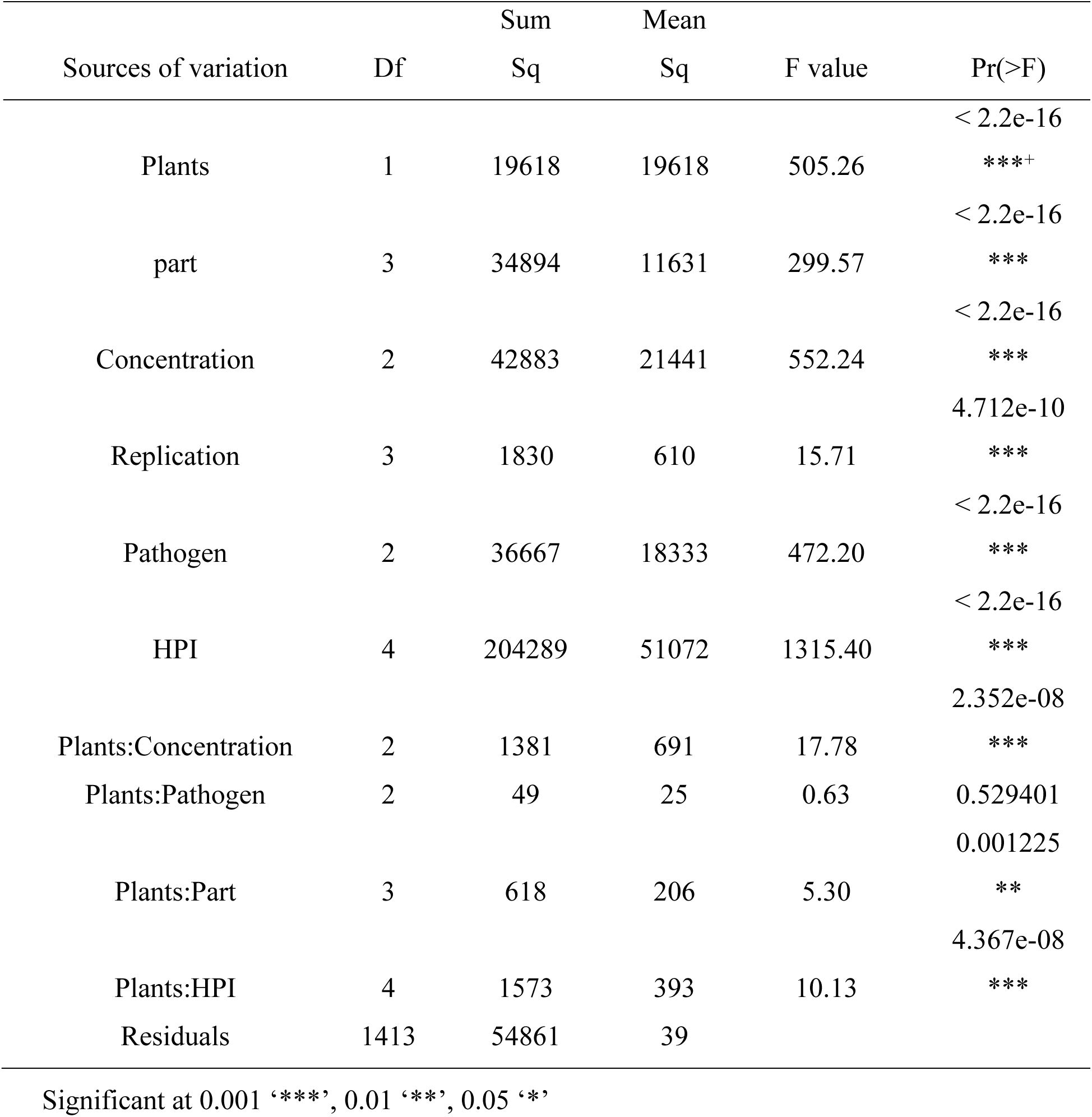
Analysis of variance of percentage of mortality.

**Figure 1:**
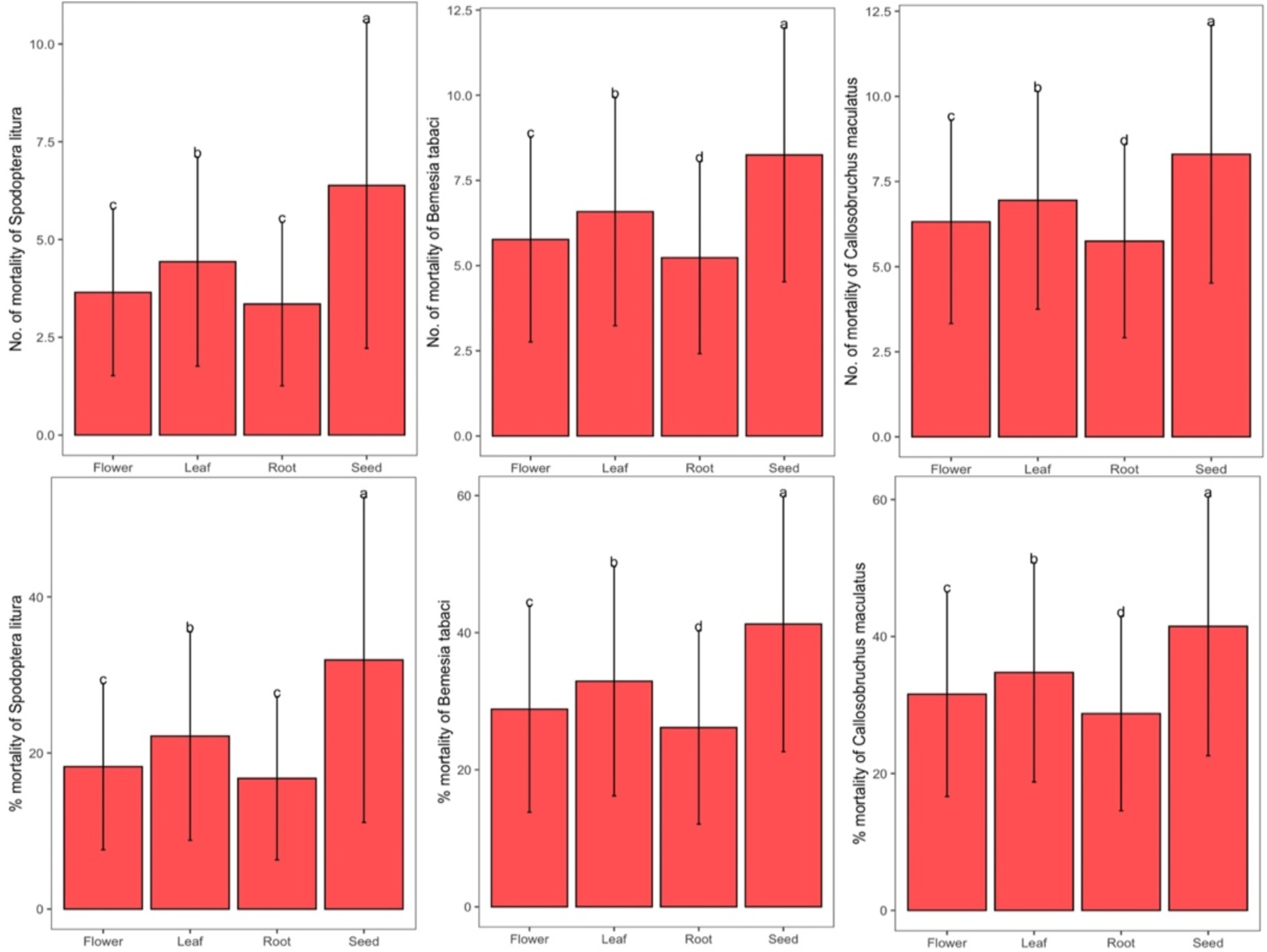
Post-hock test (LSD) (*Dhatura stramonium*) against important agricultural pests studied in this study.

**Figure 2:**
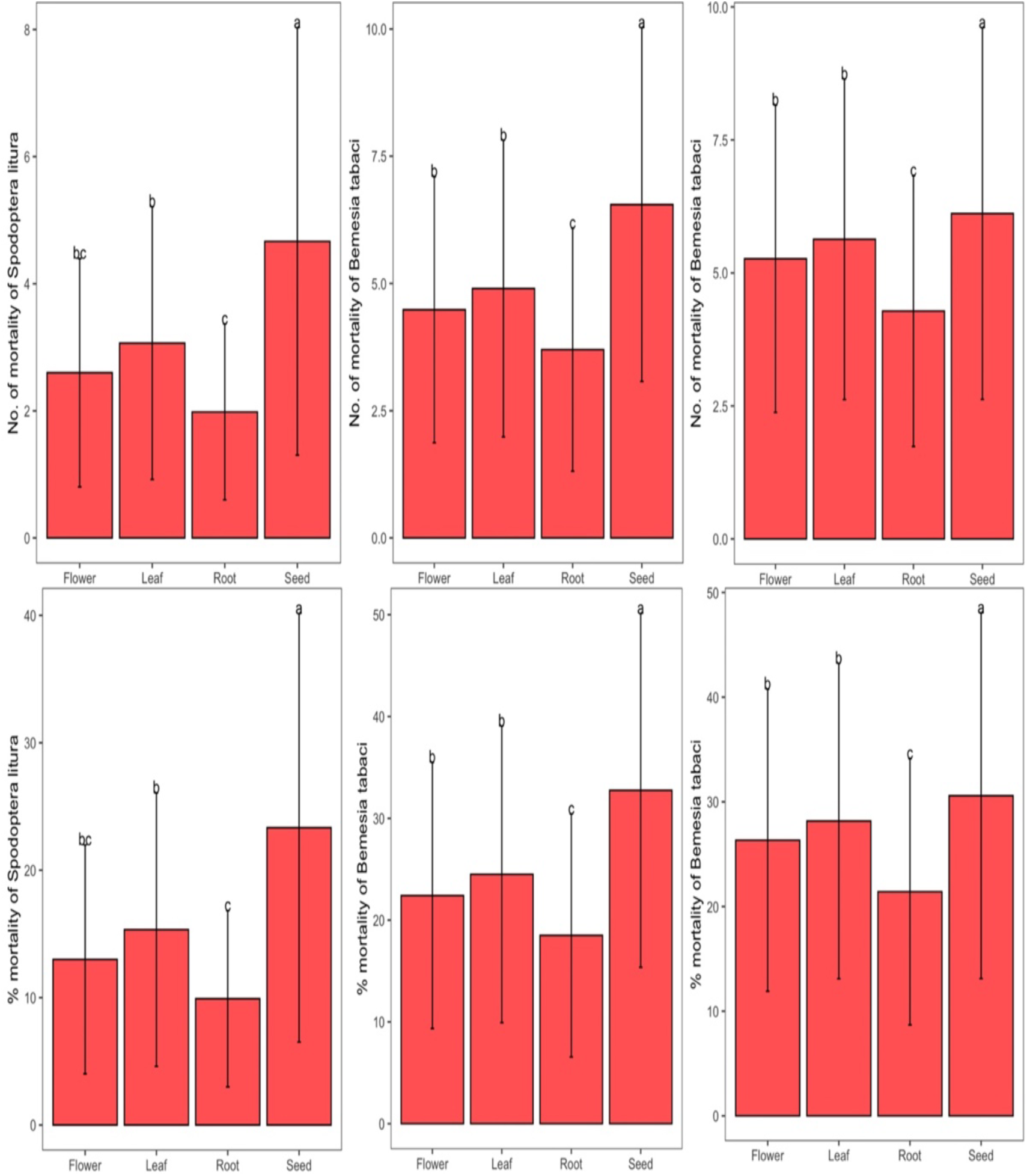
Post-hock test (LSD) (*Dhatura alba*) against important agricultural pests studied in this study.

## 4. Discussion

Plants have undergone co-evolution with insects that consume them, leading them to develop defense mechanisms. These mechanisms involve the production of compounds that interfere with insects’ usual physiological and behavioral processes. As a result, these compounds impact various aspects of insect life, such as feeding, mating, mortality, and egg-laying (Adesina, 2022). Plant-based insecticides are highly regarded as cost-effective and environmentally friendly options for safeguarding plants and stored grains. Numerous plant extracts have been employed for the management of different insect pests that infest stored goods. The essential oils derived from aromatic plants have gained recognition for their diverse properties, including cytotoxicity, antioxidant activity, antifungal effects, insecticidal properties, and antibacterial capabilities (Zimmermann et al, 2021). In studies involving direct contact and spray application, leaf extracts of *Datura metel* (extracted using acetone, water, and petroleum ether) have demonstrated insecticidal and insect repellent properties against a range of insect species. Similarly, in the case of *Datura stramonium*, the pesticide activity of non-polar extracts has been evaluated against adult insects and larvae through both contact and ingestion (Maheshwari et al, 2013).

The current study examined the effectiveness of *Datura stramonium* and *Datura alba* extracts against three significant pests: *Spodoptera litura*, *Bemesia tabaci*, and *Callosobruchus maculatus*. Among these pests, *Callosobruchus maculatus* is particularly detrimental as it causes more than 90% of damage to stored grains within a short time frame. Through bioassays, the researchers evaluated the mortality rates of these pests when exposed to extracts from the roots, leaves, flowers, and seeds of *Datura stramonium* and *Datura alba*.

The statistical analysis result after conducting ANOVA for all the traits examined in the study revealed that there are significant variations attributed to various factors, including plants, plant parts, concentration, replication, pathogens, hours post-inoculation (HPI), and their interactions. Notably, the interactions between plants and concentration, plants and pathogens, plants and plant parts, and plants and HPI were found to be highly significant. These findings were anticipated given the complex nature of phenotyping involved in the study. Furthermore, when the mortality rates of the larvae at different concentrations and over different time intervals were examined, it was found that the mortality percentage of larvae of these pests was dependent on both the concentration and duration of exposure. The mortality percentage of *Spodoptera litura* larvae was concentration and time-dependent. Notably, higher concentrations were associated with increased mortality. Also, the effectiveness of the extracts varied across different plant parts and concentrations.

Earlier, Shagal et al. (2012) conducted a study to investigate the synthesis of hyoscyamine and scopolamine in different plant parts and developmental stages of *Datura stramonium*.

While the entire plant possesses toxicity, the research by Shagal et al. (2012) and Oseni et al. (2011) revealed that the mature seeds contain the highest concentration of alkaloids. Similar to the prior investigations, the present results also showed that the mortality rates of all three pests were high when they were subjected to the seeds of the *Datura stramonium* and *Datura alba* followed by the leaf flower and root respectively. Higher concentrations generally correlated with increased mortality as the mortality rates of *Bemesia tabaci* larvae were also concentration and time-dependent. In addition, longer exposure times consistently resulted in higher mortality rates. Similarly, concentration-dependent mortality was also observed for *Callosobruchus maculatus*.

The efficacy of the extracts varied based on the specific plant part and concentration used. Studies by Taha and Mahdi (1984), Diker et al. (2007), and Boumba et al. (2005) demonstrated that ethanol extracts derived from *D. stramonium* leaves and seeds exhibit acaricidal, repellent, and oviposition deterrent effects against spider mites (*T. urticae*). Previous research has documented that when D. stramonium aqueous root extract was tested on two mosquito species, it displayed larvicidal efficacy ranging from 50% to 100% within 24 hours of treatment, employing a 100% concentration of the extracts (Singh et al, 2023).

In a separate investigation, methanol extracts from *D. stramonium* and *D. inoxia* were found to have a dose-dependent activity against Gram-positive bacteria, but their impact on *Escherichia coli* and *Pseudomonas aeruginosa* was limited. Furthermore, Takhi and Ouinten (2011) and Sharma and Sharma (2010) evaluated the combined crude ethanolic extract of *D. stramonium*, *Terminalia arjuna*, and *Withania somnifera* for its antibacterial and antifungal properties using the cup plate diffusion method. In another study conducted by Khandare and Salve, 2011, it was confirmed that leaf extracts obtained from *D. stramonium* exhibited antifungal properties against *Fusarium oxysporum*, a fungal pathogen causing wilt in pigeon pea (*Cajanus cajan* L.). Research has shown the effectiveness of different concentrations of an aqueous extract obtained from *D. stramonium* leaves and seeds in controlling flea beetles, prevalent pests in maize (Sakadzo et al, 2018).

Moreover, surviving test subjects exposed to the toxic impact of *Datura inoxia* acetone extracts exhibited inhibition of enzymes, including acetylcholinesterase, carboxylesterase, acid phosphatases, and alkaline phosphatases (ALP) when tested against *Tribolium castaneum, Trogoderma granarium*, and Sitophilus granaries (Sakadzo et al, 2018). Experiments conducted both in vivo and in vitro revealed that elevated concentrations of ethanolic leaf extracts led to the total inhibition of linear growth and sporulation in the tested fungi.

This study has unveiled a correlation between the concentration of the insect larvae and their mortality, with higher concentrations resulting in elevated mortality rates. In summary, the results suggest that the insecticidal efficacy of Datura plant extracts is influenced by factors such as the source material, concentration, and exposure duration. Extended exposure, particularly over 72 hours, consistently heightened mortality rates. These findings offer valuable insights into the potential utilization of Datura plant extracts for agricultural insect pest control.

Botanical solutions are resurgent in the realm of biopesticides thanks to their eco-friendly attributes. Plants provide a bountiful supply of bioactive compounds, rendering them promising substitutes for traditional insect-control agents. The investigation of secondary metabolites within the *Datura* genus has demonstrated significant relevance and has resulted in vital biological discoveries. Constant research endeavours are uncovering novel metabolites that demonstrate potential biological effects in diverse systems. As a result, the *Datura* genus is acknowledged as a valuable source of chemical compounds with distinct pharmaceutical uses, and one particularly captivating feature is its effectiveness in controlling insect populations.

## 5. Conclusion

The study revealed the significant insecticidal potential of *Datura* plant extracts derived from various plant parts, including roots, leaves, flowers, and seeds, against three important pests: *Spodoptera litura, Bemesia tabaci,* and *Callosobruchus maculatus*. Notably, higher concentrations of Datura plant extracts were linked to increased mortality rates across all three pest species, highlighting the critical need for optimizing extract concentrations to achieve effective pest control. Furthermore, the research unveiled the time-dependent efficacy of Datura plant extracts. Extended exposure, especially beyond 72 hours, consistently led to higher mortality rates, emphasizing the importance of considering exposure duration when devising pest control strategies.

The efficacy of Datura plant extracts exhibited variations depending on the specific plant part used, with leaf extracts, particularly at higher concentrations, demonstrating remarkable insecticidal effects. These findings provide valuable guidance for selecting the most suitable plant components for extract preparation. The comprehensive statistical analysis underscored the intricate nature of phenotyping within this study. Multiple factors, including the type of plants, plant components, extract concentration, replication, pathogens, hours post-inoculation (HPI), and their interactions, all played highly significant roles in shaping the experimental outcomes. This highlights the multifaceted nature of investigating the insecticidal properties of these extracts. Crucially, these findings have practical implications for developing environmentally friendly and efficient pest control strategies in agriculture. By carefully considering extract concentration, exposure duration, and the choice of plant components in Datura plant extract formulations, tailored approaches can be devised to address specific pest management needs.

## Notes

### Competing Interest Statement

The authors have declared no competing interest.

## Reference

1. Hsiao TH, Fraenkel G (1968) The role of secondary plant substances in the food specificity of the Colorado potato beetle. Ann Entomol Soc Am 61:485–503

2. Krug E, Proksch P (1993) Influence of dietary alkaloids on survival and growth of Spodoptera littoralis. Biochem Syst Ecol 21:749–756

3. Kuganathan N, Ganeshalingam S (2011) Chemical analysis of Datura metel leaves and investigation of the acute toxicity on grasshoppers and red ants. E-J Chem 8:107–112 Kuganathan N, Saminathan S, Muttukrishna S (2008) Toxicity of Datura alba leaf extract to aphids and ants. Internet J Toxicol 5:2

4. Zhou J, Hu GF, Liu MY, Li YQ, Li JJ, Yu HT (2008) Contact and antifeeding effects on four species of the genus Datura against Mythimna separate and Pieris rapae. J Gansu Agric Univ 43:102–106

5. Malik MF (2005) Bionomic field studies and integrated pest management of thrips with species innovation through agroecosystem of onion (Allium cepa) in Balochistan Pakistan. Dissertation, University of Balochistan, Quetta

6. Bourgaud F, Gravot A, Milesi S, Gontier E (2001) Production of plant secondary metabolites: a historical perspective. Plant Sci 161:839–851

7. Gerlach GH (2006) Datura innoxia-a potential commercial source of scopolamine. Econ Bot 2:436–454

8. Shonle I, Bergelson J (2000) Evolutionary ecology of the Tropane alkaloids of Datura stramonium L. (Solanaceae). Evolution 54:778–788

9. Belmain, S. R., Neal, G. E., Ray, D. E. and Golop, P. 2001: Insecticidal and vertebrate toxicity associated with ethnobotanicals used as postharvest protectants in Ghana. Food Chem. Toxicol. 39: 287–291.

10. Bayih T (2014) Synergistic bio-eficacy of insecticidal plants against bean bruchids (Zabrotessubfasciatus: Coleoptera) a major storage pests of common bean (Phaseolus vulgaris L.) in central rift valley of Ethiopia. MSc thesis, Department of Biology, School of Graduate Studies, Haramaya University.

11. Shaaya, E., Kostjukovski, M., Eilberg, J. and Sukprakarm, C. 1997: Plant oils as fumigants and contact insecticides for the control of stored product insects. J. Stored Prod. Res. 33: 7–15.

12. Ho, S. H., Ma, Y. and Huang, Y. 1997: Anethole, a potential insecticide from Illicium verum Hook F., against two stored-product insects. Int. Pest Control 39: 50–51.

13. Huang, Y., Tan, J. M. W. L., Kini, R. M. and Ho, S. H. 1997: Toxicity and antifeedant action of nutmeg oil against T. castaneum (Herbst) and Sitophilus zeamais Motsch. J. Stored Prod. Res. 33: 289–298.

14. Huang, Y., Lam, S. L. and Ho, S. H. 2000: Bioactivities of essential oil from Elletaria cardamomum (L.) Maton. to Sitophilus zeamais Motschulsky and Tribolium castaneum (Herbst). J. Stored Prod. Res. 36: 107–117.

15. Dev, S. and Koul, O. 1997: Insecticides of natural origin. New York: Harwood Academic, 352 pp.

16. Kim, S., Roh, J. Y., Kim, D. H., Lee, H. S. and Ahn, Y. J. 2003: Insecticidal activities of aromatic plant extracts and essential oils against Sitophilus oryzae and Callosobruchus chinensis. J. Stored Prod. Res. 39: 293–303.

17. Saxena, R. C., Dixit, O. P. and Sukumaran, P. 1992: Laboratory assessment of indigenous plant extracts for antijuvenile hormone activity in Culex quinquefasciatum. Indian J. Med. Res. 95: 204–206.

18. Shaalan, E. A. S., Canyon, D., Younes, M. W. F., Abdelwahab, H. and Mansour, A. H. 2005: A review of botanical phytochemicals with mosquitocidal potential. Environ. Intern. 31: 1149–1166.

19. Soni P, Siddiqui AA, Dwivedi J, Soni V (2012) Pharmacological properties of Datura stramonium L. asa potential medicinal tree: An overview. Asian Pac J Trop Biomed 2(12):1002–1008.

20. Li J, Lin B, Wang G, Gao H Qin M (2012) Chemical constituents of Datura stramonium seeds. China Journal of Chinese material 37(3): 319–322. Usha K, Singh B, Praseetha P, Deepa N, Agarwal DK, et al. (2009) Antifungal activity of Datura stramonium, Calotropisgigantea and Azadirachtaindica against Fusarium mangiferae and malformation in mango, European Journal of Plant Pathology 124: 637–657.

21. Olofintoye LK, Simon IA, Omoregie OB (2011) Larvicidal properties of Datura stramonium (Jimson weed) and Nicotianatabaccum extracts against the larvae of (Anopheles and Culex) mosquitoes. African research review V 5(2): 337–344.

22. (2000) PDR for Herbal Medicines. Medical Economics Company, Inc. at Montvale pp. 436–437.

23. Tostes RA (2002) Accidental Datura stramonium poisoning in a dog. Veterinary and Human Toxicology 44(1): 33–34.

24. Priya KA, Gnanamani A, Radhakrishnan N, Babu M (2002) Healing potential of Datura alba on burn wounds in albino rats. J Ethnopharmacol 83:193–199

25. Evans W (1979) Tropane alkaloids of the Solanaceae. In: Hawkes JG, Lester RN, Skelding A (eds) The biology and taxonomy of the Solanaceae. Academic Press, London, pp 241–254

26. Conklin M (1976) Genetic and biochemical aspects of the development of Datura. In: Wolsky E (ed) Monographs in developmental biology, vol 12. S. Karger, New York, p 170.

27. S.-T. Fang, X. Liu, N.-N. Kong, S.-J. Liu, and C.-H. Xia, “Two new withanolides from the halophyte Datura stramonium L.,” Natural Product Research (Formerly Natural Product Letters), vol. 27, no. 21, pp. 1965–1970, 2013

28. Khandare KR, Salve SB. Management of wilt of pigeon pea (*Cajanus cajan* L.) through biopesticide (leaf extracts) Int Refer Res J. 2011;2(18):21–22.

29. Adesina, J.M. Bioactive constituents and fumigant toxicity of Datura metel extracts as grain protectant and progeny emergence inhibition of *Callosobruchus maculatus* (Coleoptera: Bruchidae). J. Plant Dis. Prot. 2022, 129, 819–829.

30. Taha SA, Mahdi AW. Datura intoxication in Riyadh. Trans R Soc Trop Med Hgy. 1984;78:134–135

31. Diker D, Markovitz D, Rothman M, Sendovski U. Coma as a presenting sign of *Datura stramonium* seed tea poisoning. Eur J Int Med. 2007;18(4):336–338

32. Boumba A, Mitselou A, Vougiouklakis T. Fatal poisoning from ingestion of *Datura stramonium* seeds. Vet Human Toxicol. 2005;46:81–82.

33. Takhi D, Ouinten M. Study of antimicrobial activity of secondary metabolites extracted from spontaneous plants from the area of Laghouat, Algeria. Adv Environm Biol. 2011;5(2):469–476.

34. Sharma MC, Sharma S. Phytochemical, preliminary pharmacognostical and antimicrobial evaluation of combined crude aqueous extract. Int J Microbiol Res. 2010;1(3):166–170.

35. Shagal MH, Modibbo UU, Liman AB. Pharmacological justification for the ethnomedical use of *Datura stramonium* stem-bark extract in treatment of diseases caused by some pathogenic bacteria. Int Res Pharm Pharmaco. 2012;2(1):16–19.

36. Oseni OA, Olarinoye CO, Amoo IA. Studies on chemical compositions and functional properties of thorn apple (*Datura stramonium* L) Solanaceae. Afric J Food Sci. 2011;5(2):40–44.

37. Zimmermann, R.C.; de Carvalho Aragao, C.E.; de Araújo, P.J.P.; Benatto, A.; Chaaban, A.; Martins, C.E.N.; do Amaral, W.;Cipriano, R.R.; Zawadneak, M.A. Insecticide activity and toxicity of essential oils against two stored-product insects. Crop Prot. 2021, 144, 105575.

38. Maheshwari NO, Khan A, Chopade BA. Rediscovering the medicinal properties of Datura sp.: A review. Journal of Medicinal Plants Research. 2013;7(39):2885–2897

39. Benhamou N, Lafontaine PJ, Nicole M. Induction of Systemic Resistance to Fusarium Crown and Root Rot in Tomato Plants by Seed Treatment with Chitosan. American Phytopath. Society, 2012; 84(12):1432–44.

40. European Commission. EU Pesticides database. ec.europa.eu. Retrieved, 2020.

41. Singh, S. and Kumar, S., 2021. Medicinal plant sector in India: Status and sustainability. International Journal of Economic Plants 8,81–85.

42. Iyekowa, O., Uwumarongie, O.H. and Okunzuwa, I.G., 2023. High-Performance Liquid Chromatography (HPLC) profile and anxiolytic activity of hexane extract of Datura metel leaves extract in balb/c mice. Journal of Science and Technology Research 5.

43. Jabeen, N., Khan, I.H. and Javaid, A., 2022. Fungicidal potential of leaf extract of Datura metel L. to control Sclerotium rolfsii Sacc. Allelopathy J 56,59–68.

44. Singh, S., Minj, K.H., Devhare, L.D., Uppalwar, S.V., Anand, S., Suman, A. and Devhare, D.L., 2023. An update on morphology, mechanism, lethality, and management of dhatura poisoning. Eur. Chem. Bull, 12(5), pp.3418–3426.

45. Sakadzo N, Pahla I, Muzemu S, Mandumbu R, Makaza K. Herbicidal effects of Datura stramonium (L.) leaf extracts on Amaranthus hybridus (L.) and Tagetes minuta (L.). African Journal of Agricultural Research. 2018; 13 (34):1754–1760

46. Hu S, Wang XY, Yang ZQ, Duan JJ. Effects of photoperiod and light intensity on wing dimorphism and development in the parasitoid Sclerodermus pupariae (Hymenoptera: Bethylidae). Biological Control. 2019 Jun 1;133:117–22.

47. Harve, G. and Kamath, V., 2004. Larvicidal activity of plant extracts used alone and in combination with known synthetic larvicidal agents against Aedes aegypti.

48. Mallick S, Mukherjee D, Chandra G. Evaluation of larvicidal efficacy of acetone leaf extracts of Annona reticulata Linn. against Aedes aegypti, Anopheles stephensi and Culex quinquefasciatus (Diptera: Culicidae). Journal of mosquito research. 2015 Jun 23;5.

49. Ali W, Annon MR. Biological Effective of organic solvent extracts of Mirabilis jalapa Leaves in the Non-cumulative for mortality of Immaturestages Culex quinquefasciatus Say (Diptera: Culicidae). Al-Qadisiyah Journal Of Pure Science. 2020;25(1):1–6.

50. Finney DJ. Probit analysis, Cambridge University Press. Cambridge, UK. 1971.

